# DNA-damaging combination treatments impose genotype-specific constraints on hypermutator evolvability

**DOI:** 10.64898/2026.02.20.706947

**Authors:** Adam J. Mulkern, Stefan O. Bassler, William Matlock, Athanasios Typas, R. Craig MacLean

**Author notes:** These authors contributed equally (Adam J. Mulkern, Stefan O. Bassler).

## Abstract

Bacterial hypermutator strains drive rapid evolution of antibiotic resistance in chronic infections. Inspired by cancer therapy approaches that exploit synthetic lethality by targeting DNA repair deficiencies in hypermutator tumours, we tested whether pairing a conventional antibiotic with a secondary DNA-damaging agent could constrain hypermutator evolution in bacteria. Using high-throughput experimental evolution of *Escherichia coli* repair-deficient strains, we evolved populations under carbapenem selection in combination with ciprofloxacin or mitomycin C. Strains lacking oxidative damage repair, double-strand break repair, or transcription-coupled repair showed significantly reduced evolvability, particularly under constant antibiotic pressure and increasing genotoxic stress. However, mismatch repair (MMR) hypermutators, the predominant clinical genotype, did not show reduced evolvability under these combination treatments. This is consistent with pathway orthogonality: MMR does not repair the structural DNA lesions induced by ciprofloxacin or mitomycin C, and the elevated mutation supply of MMR-deficient strains may allow rapid adaptation despite background DNA damage. Our findings demonstrate that combination strategies can constrain the evolvability of specific repair-deficient genotypes *in vitro*, but success requires matching DNA damage type to specific repair vulnerabilities. This work establishes proof of principle for genotype-directed antimicrobial strategies that exploit DNA repair vulnerabilities to constrain hypermutator evolution.

**Significance:** This work demonstrates that combining DNA-damaging agents with antibiotics can constrain resistance evolution in specific hypermutator genotypes, but not others. These findings establish that evolution-informed antimicrobial strategies must be genotype-specific, opening new paths for precision approaches to delay or prevent the evolution of antibiotic resistance by exploiting DNA repair vulnerabilities.

## Introduction

Antimicrobial resistance (AMR) is a critical global issue, creating an urgent need to develop rationally designed treatment strategies to outpace the evolution of resistance^1^. Of particular interest in this context are bacterial hypermutator strains with elevated mutation rates due to defects in DNA repair^2^. These strains, though rare in most natural populations, are relatively frequent in chronic infections that occur in patients with cystic fibrosis, where they drive resistance evolution^3,4^. Consequently, understanding these strains is urgent, as they can rapidly evolve resistance to both antibiotics and bacteriophages^5^, often acquiring compensatory adaptations that offset the biological cost of resistance^6^.

In parallel, analogous evolutionary dynamics are observed in oncology, where hypermutation allows tumours to acquire resistance to chemotherapeutic drugs^7^. The underlying defects in DNA repair found in hypermutating tumours create a state of genomic instability that can be exploited therapeutically by treating tumours with DNA-damaging agents^8^. For example, BRCA1/BRCA2-deficient breast and ovarian tumours with impaired homologous recombination show synthetic lethality when combined with inhibitors of the single-strand-break repair enzyme poly(ADP-ribose) polymerase 1 (PARP1)^9^. Drawing inspiration from cancer therapies that exploit such vulnerabilities^10^, we aim to determine whether targeting DNA repair-deficient bacterial strains with DNA-damaging agents can exploit synthetic lethal interactions to selectively eliminate hypermutator lineages. By specifically targeting hypermutating genotypes, this approach could potentially suppress the emergence of resistant subpopulations, offering a novel, evolution-informed strategy to delay or prevent AMR evolution.

To investigate whether the principle of synthetic lethality can be translated from oncology to bacterial AMR, we used high-throughput experimental evolution of a panel of *Escherichia coli* knockout mutants from the KEIO library^11,12^. To map which DNA repair mechanisms influence evolutionary trajectories, we included gene deletions of mismatch repair (Δ*mutS*, Δ*mutH*, and Δ*mutL*), oxidative damage repair (Δ*mutM*, Δ*mutT*, and Δ*mutY*), double-strand break repair (Δ*recA* and Δ*recN*) and transcription-coupled repair (Δ*mfd*). We subjected these strains to diverse selective regimes, coupling the carbapenem antibiotic meropenem with DNA-damaging agents (ciprofloxacin and mitomycin C) delivered at either constant sub-inhibitory levels or increasing concentrations. By quantifying growth dynamics across these regimens, we sought to determine if exacerbating genomic instability could effectively constrain the evolvability of hypermutator strains and suppress the emergence of resistance.

## Results

Using the *Escherichia coli* K-12 (BW25113) KEIO gene-knockout collection^11,12^, we evolved classical mutator strains carrying gene deletions in *mutS*, *mutH*, and *mutL* (mismatch repair) and in *mutM*, *mutT*, and *mutY* (oxidative damage repair). We also evolved strains carrying gene deletions in *recA* and *recN* (homologous recombination/double-strand break repair) and *mfd* (transcription-coupled repair) to broaden the DNA-repair landscape beyond mutation avoidance alone.

To test the hypothesis that pairing antibiotic selection with targeted DNA damage can constrain the evolution of hypermutators, we measured the ability of bacterial populations to adapt to increasing antibiotic doses, with or without a second bactericidal agent that induces DNA damage. We selected meropenem as the primary antibiotic selective pressure to drive resistance evolution because of its clinical relevance. To induce the requisite replication stress, we selected two distinct DNA-damaging agents to pair with it: ciprofloxacin, a fluoroquinolone that induces double-strand breaks via topoisomerase inhibition^13^, and mitomycin C, which causes inter-strand DNA crosslinks^14^.

We used high-throughput *in vitro* laboratory evolution with a stepwise ‘drug selection ramp’ to drive adaptation^15,16^. All strains were evolved under defined drug selection as follows: populations were passaged daily by 1:100 dilution (corresponding to a bottleneck of ~10⁶ cells per transfer at saturation density) under two-fold daily increases from 0.5 x MIC to 32 x MIC, where MIC values were determined for the wildtype strain (meropenem = 0.125 µg/mL, ciprofloxacin = 0.016 µg/mL, mitomycin C = 1 µg/mL). For combination treatment regimes, the focal drug followed the same ramp while the partner drug was maintained at a fixed sub-inhibitory concentration throughout (**Fig. 1**).

**Figure 1.**
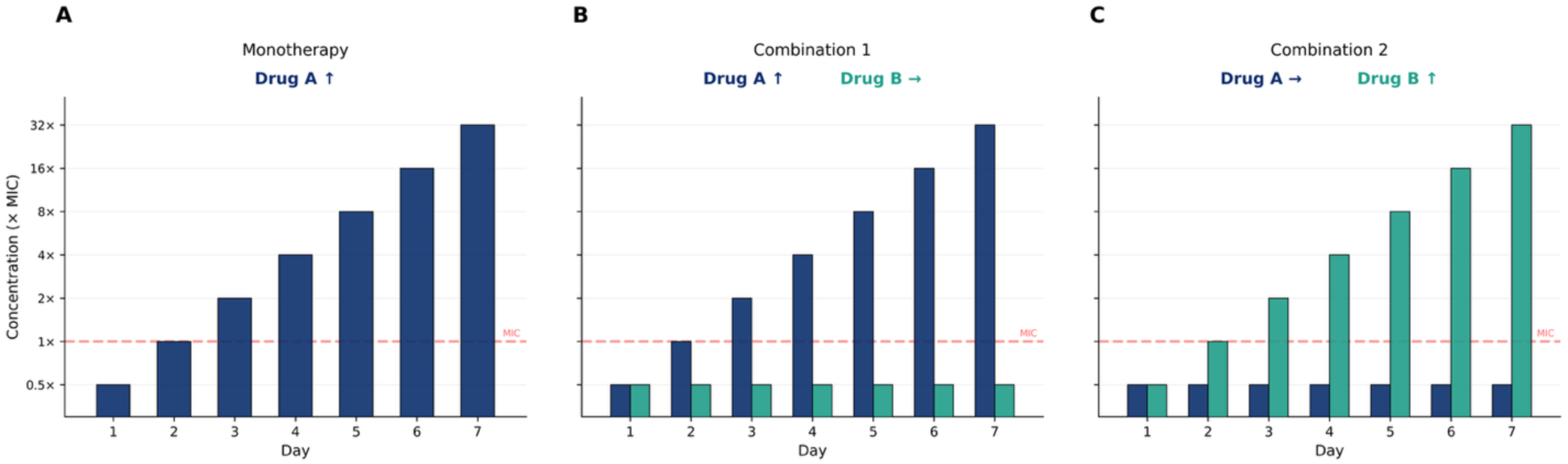
Experimental design. Stepwise drug selection ramp over seven days of experimental evolution, populations were passaged daily at a 1:100 dilution. (A) Monotherapy: the focal drug concentration doubles daily from 0.5× to 32× MIC. (B) Combination (Drug A ↑ Drug B →): Drug A follows the escalating ramp while Drug B is maintained at a constant sub-inhibitory concentration (0.5× MIC). (C) Combination (Drug A → Drug B ↑): the inverse orientation. Both combination orientations were tested for each antibiotic-DNA-damaging agent pair.

### Growth dynamics uncover the emergence of selective constraints

We first examined the aggregate evolutionary trajectories of these genotypes by grouping them into four functional classes: Mismatch Repair (MMR), Oxidative Damage Repair (ODR), Recombination/Double Strand Break Repair (RR), and Transcription-Coupled Repair (TCR) (**Fig. 2**). This high-level overview revealed distinct patterns of growth. While the MMR-deficient populations (Δ*mutS*, Δ*mutH*, Δ*mutL*) generally maintained high optical densities throughout the dose increase, the Recombination (Δ*recA*, Δ*recN*) and Transcription-Coupled Repair (Δ*mfd*) groups showed rapid population collapse, particularly in regimes with ciprofloxacin. The ODR group (Δ*mutM*, Δ*mutT*, Δ*mutY*) displayed intermediate phenotypes, often diverging from the WT baseline in specific combination treatments (e.g., Mero → Mito ↑), indicating a regime-dependent vulnerability.

**Figure 2.**
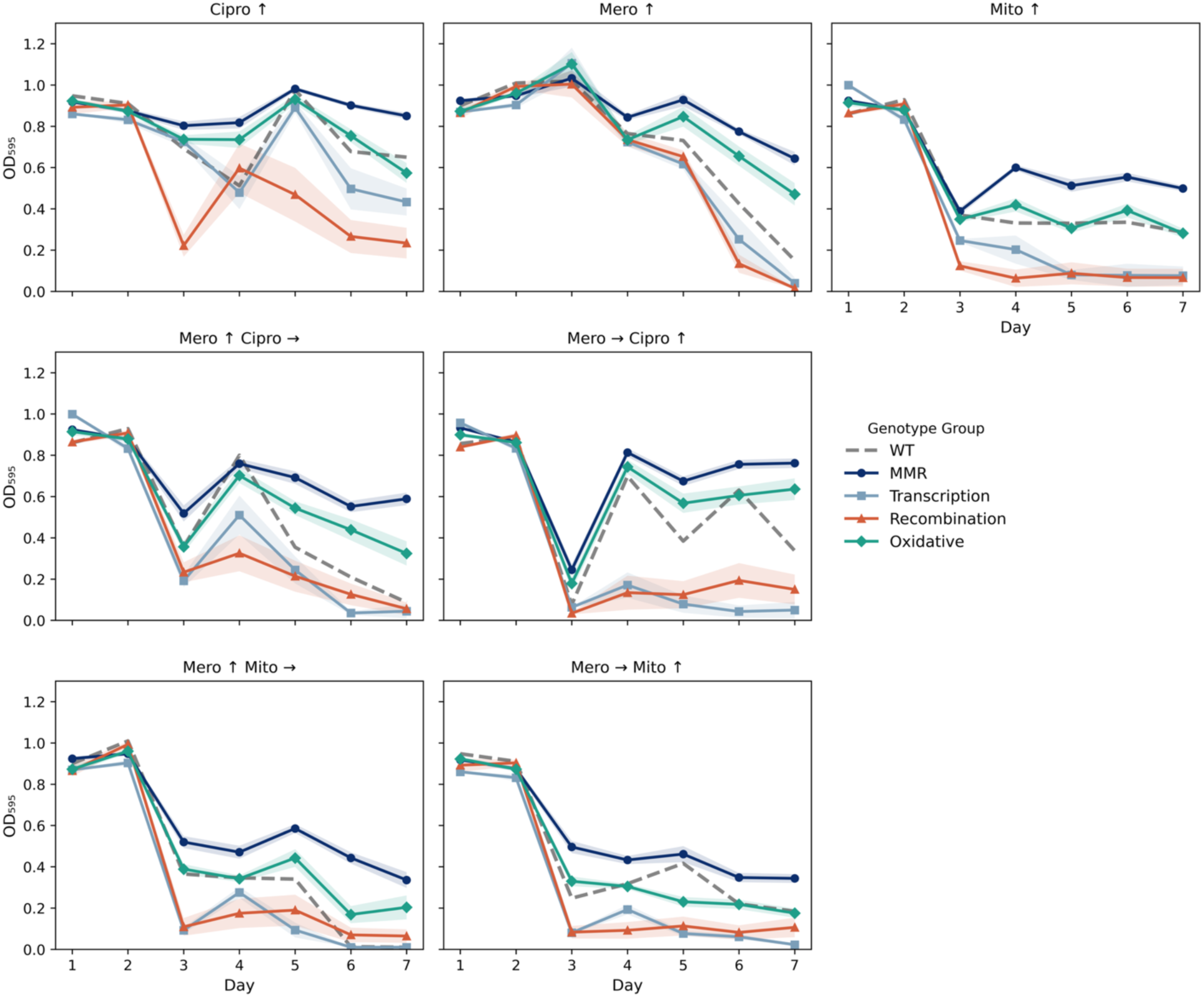
Functional Group Growth Dynamics. Mean optical density (OD_595_) trajectories are shown over 7 days of experimental evolution, with OD measured every 24 h prior to transfer. Genotypes are aggregated into four functional classes: Mismatch Repair (MMR: Δ*mutH*, Δ*mutL*, Δ*mutS*; n = 24), Transcription-Coupled Repair (TCR: Δ*mfd*; n = 8), Recombination/DSB Repair (RR: Δ*recA*, *ΔrecN;* n = 16), and Oxidative Damage Repair (ODR: Δ*mutM*, Δ*mutT*, Δ*mutY*; n = 24). WT is shown as a dashed grey line (n = 8). Shaded regions represent ± s.e.m. calculated across all biological replicates and strains within each functional group.

We assessed evolutionary trajectories by tracking the growth difference between each mutant and its wild-type (WT) control (ΔΔOD_595 nm_ = OD_mutant 595 nm_ – OD_wt control 595 nm_) for every genotype, drug regime, and day (**Fig. 3**). Starting concentrations of the antibiotics were selected to not inhibit growth alone and in combination to start with comparable population sizes, which can be seen at similar OD during the initial phase of the experiment (Day 1 and Day 2), across treatments and genotypes. This suggests that the baseline susceptibility of the mutants to the starting antibiotic concentrations is comparable to that of the WT.

**Figure 3.**
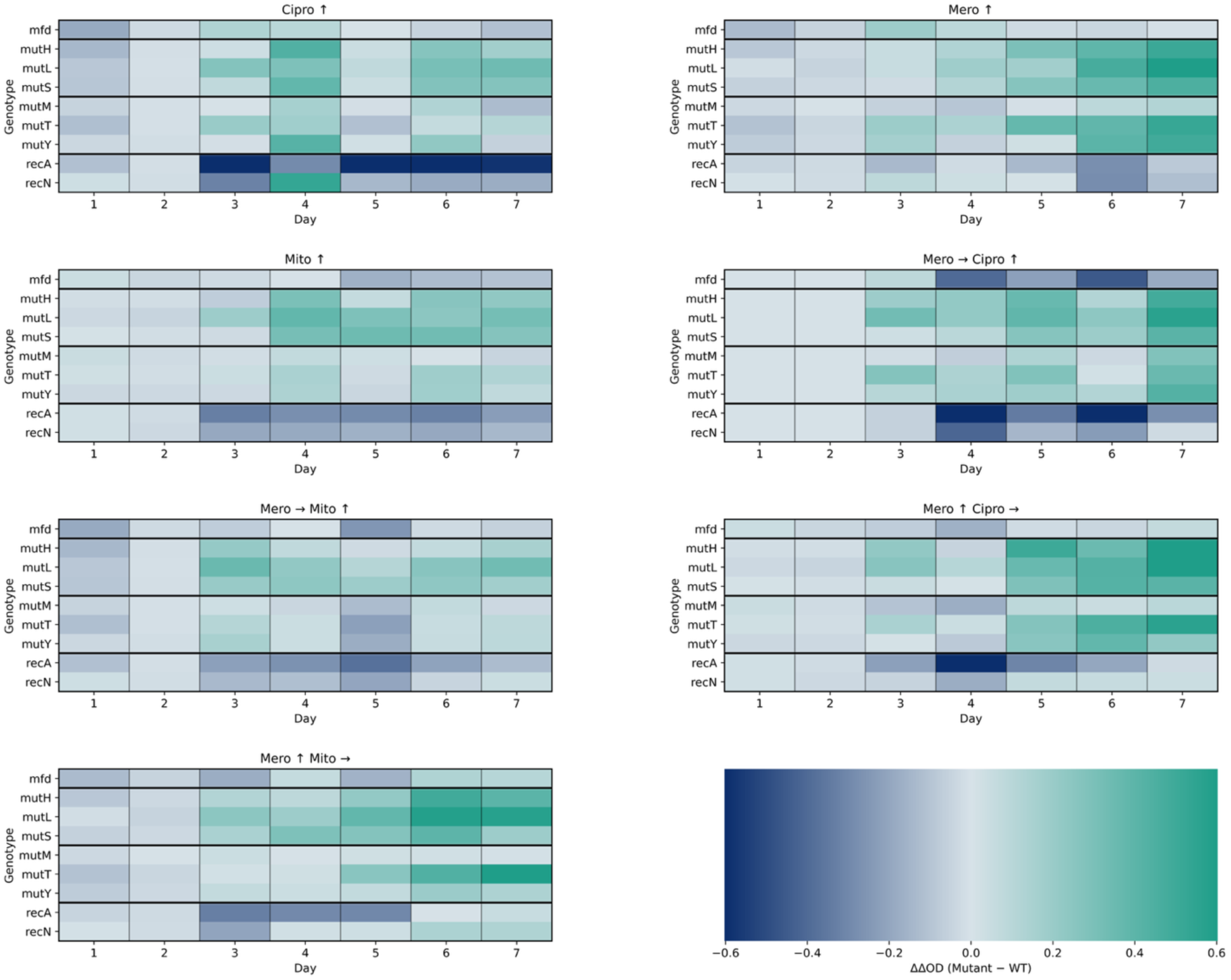
Growth Dynamics. Background-subtracted growth of mutant relative to WT over time. Heatmaps show ΔΔOD_595 nm_= OD_mutant 595 nm_ – OD_wt 595 nm_ for each genotype–treatment–day combination, values are averaged across replicates. Colours use a diverging scale centered at 0, with extremes clipped at the 98th percentile to preserve contrast: green (>0) indicates higher OD than WT (reduced inhibition), grey (≈0) indicates no difference, and blue (<0) indicates lower OD than WT (greater inhibition). Standard growth curves are shown in **Supplementary Fig. 1**.

As drug concentrations increased, populations could only persist by adapting to the treatment regime. Divergence between mutants and the WT emerged from Day 3 onwards, coinciding with the point at which the selection ramp breached the initial MIC threshold. As anticipated, control genotypes deficient in essential DNA repair and stress response pathways (Δ*recA*, Δ*recN*, and Δ*mfd*) displayed sustained negative ΔΔOD values relative to the WT from Day 3 onwards. This confirmed that our experimental regime successfully imposed selective suppression on strains unable to tolerate DNA replication stress (**Supplementary Fig. 1**). However, in hypermutator genotypes, the dynamics were more complex, with divergent trajectories emerging between Days 5 and 7 depending on the specific repair defect.

### Quantifying differences in evolutionary potential

To formally quantify these differences in evolvability, we analysed area-under-the-curve (AUC) measurements from mutant and wildtype bacterial growth using a Bayesian model. The key parameter of interest was the mutant–treatment effect, which represented the typical difference in AUC (ΔAUC) between mutant and wildtype under a given treatment (**Fig. 4**). The model is described in full in **Supplementary File 1**.

**Figure 4.**
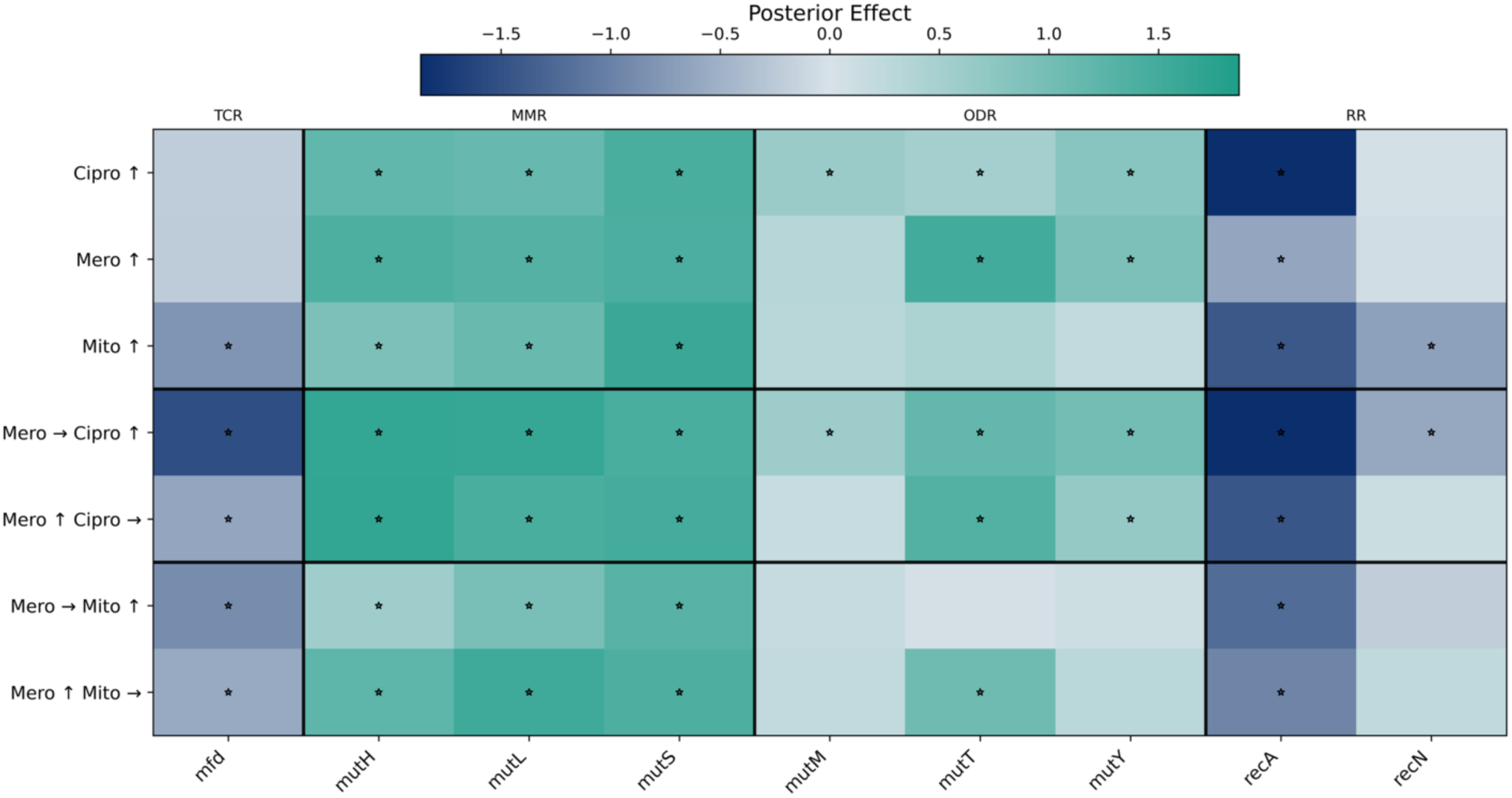
Posterior Effect. Heatmap of posterior mean ΔAUC in OD_595nm_ (Mutant − WT) for all genotype-treatment combinations, showing estimated mutant-WT growth differences across drugs and schedules. Grey (~0) denotes no difference from WT; blue (<0) indicates selective suppression of the mutant; green (>0) indicates relative mutant advantage. Asterisks (*) mark where the 95% credible interval excludes 0. Colours are centered at 0 with extremes clipped at the 98th percentile to preserve contrast. Posterior estimates derive from a Bayesian model in which WT and mutant observations are modelled as independent groups sharing drug-specific baselines, with treatment-specific shifts (ΔAUC) estimated for each mutant–drug combination (detailed in **Supplementary File 1**).

We determined that mutants with impaired double-strand break repair and transcription-coupled repair (Δ*recA*, Δ*recN*, Δ*mfd*) exhibited consistently lower evolvability than the WT control in single-drug treatments, highlighting the critical role of these pathways in adapting to antibiotic stress. The addition of DNA-damaging agents (ciprofloxacin and mitomycin C) showed no clear posterior shift, consistent with the loss of these pathways being deleterious regardless of the secondary stressor.

In contrast, the impact of defects in mutation avoidance pathways was distinct. The ODR mutants (Δ*mutM*, Δ*mutT*, Δ*mutY*) generally displayed evolvability close to zero. 11 of the 21 lineage/drug combinations showed significantly negative ΔAUC values. This indicates that while these defects do not abolish the capacity to evolve resistance, they substantially reduce evolvability relative to MMR mutants across multiple treatment contexts (**Fig. 5**). This contrasts with MMR mutants (Δ*mutS*, Δ*mutH*, Δ*mutL*), which frequently exhibited an evolutionary advantage (positive ΔAUC). Crucially, these evolutionary benefits driven by MMR-deficiency were not eradicated by the addition of the tested DNA-damaging agents in this configuration.

**Figure 5.**
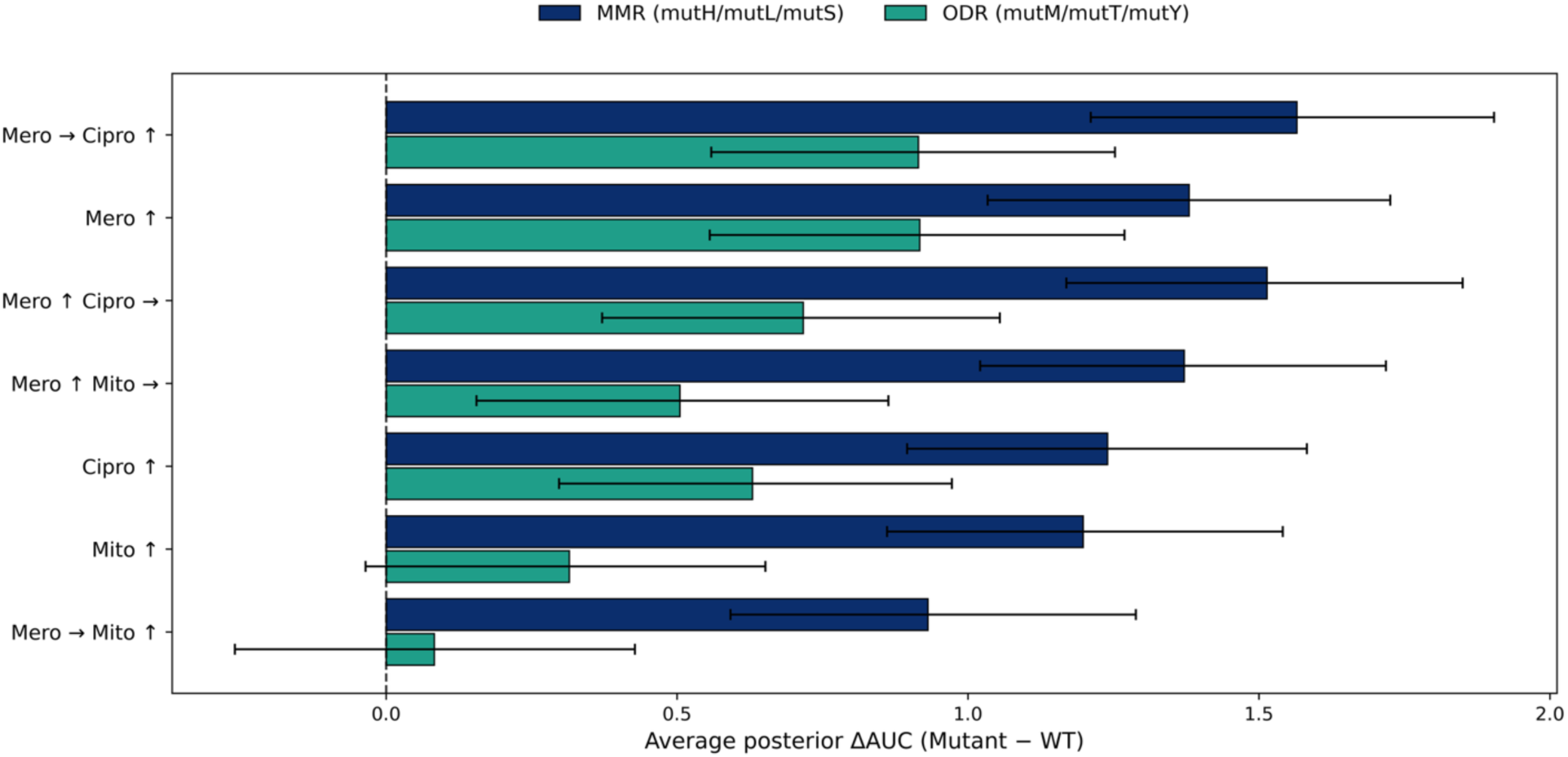
Grouped Treatment Rankings. (posterior ΔAUC, mutant − WT). Horizontal bars show, for each treatment, the average posterior mean ΔAUC (OD_595_; Mutant − WT), aggregated across mutants within each class. Blue bars correspond to MMR mutants (Δ*mutH,* Δ*mutL,* Δ*mutS*); green bars correspond to ODR (Δ*mutM,* Δ*mutT,* Δ*mutY*). Error bars indicate the 95% credible interval of the group-level posterior mean, obtained by averaging posterior draws across mutants within each class. Treatments are ordered by the grand mean across both groups (with more negative values at the top). The vertical dashed line marks parity with WT (0). Values closer to 0 indicate greater constraint on mutant evolvability relative to WT; larger positive values indicate a stronger relative mutant advantage.

### Drug regimen and orientation modulate selective constraints

While the genotype-specific trends were robust, the specific drug scheduling modulated the degree of evolutionary constraint. The regime where meropenem was held constant while mitomycin C was increased (Mero → Mito ↑) produced ΔΔOD trajectories that remained closer to zero across days for many genotypes compared with other regimens (**Fig. 3**). This indicates that when the primary antibiotic pressure is maintained at a constant concentration, mutant responses track their WT more closely, resulting in smaller divergences than observed in the inverse treatment regime where the DNA-damaging background was constant and the antibiotic was escalated (Mero ↑ Mito →).

### Oxidative damage repair mutants are broadly vulnerable to combination therapy

To synthesise the treatment effects, we pooled the posterior ΔAUC estimates for each mutant relative to its WT within two functional classes: MMR (Δ*mutH*, Δ*mutL*, Δ*mutS*) and ODR (Δ*mutM*, Δ*mutT*, Δ*mutY*) (**Fig. 5**). A distinct hierarchy of vulnerability emerged. The ODR class consistently exhibited substantially lower ΔAUC values than the MMR class across all treatments, indicating reduced evolvability of these genotypes under the tested regimens. Conversely, the MMR class remained largely robust, often retaining an evolutionary advantage despite the added genotoxic stress.

Finally, we ranked the treatment regimens by their ability to constrain these hypermutator populations. The combination regime where meropenem was held constant while mitomycin C was increased (Mero → Mito ↑) proved most efficacious at reducing mutant evolvability. Under this schedule, ODR mutants showed the smallest ΔAUC values of any treatment, approaching parity with the WT, while even MMR mutants showed their most attenuated evolutionary advantage.

### Bliss independence analysis reveals synergistic drug interactions

To determine whether combination treatments produced non-additive fitness effects, we computed Bliss independence scores for each genotype under each combination regimen. Under constant meropenem paired with increasing mitomycin C (Mero → Mito ↑), both MMR and ODR groups showed significantly lower Bliss scores than wildtype at the group level (Wilcoxon rank-sum test; MMR: Δbliss = −0.38, p < 0.0001; ODR: Δbliss = −0.23, p = 0.014), with five of six classical mutator genotypes showing individually significant synergy after Benjamini–Hochberg correction (**Supplementary Fig. 2**). Positive Bliss scores for recombination-deficient strains (ΔrecA, ΔrecN) likely reflect a floor effect rather than true antagonism, as these strains showed near-complete growth inhibition under monotherapy alone. Wildtype Bliss scores were approximately zero across all combinations (mean range: +0.001 to +0.003: all p > 0.9), serving as the reference distribution against which mutant scores were tested.

Day-resolved Bliss scores revealed the temporal dynamics of these interactions (**Fig. 6**, **Fig. 7**). Synergy emerged progressively from Day 4 onwards, coinciding with drug concentrations exceeding the MIC, and intensified through Day 7. This pattern was consistent across individual genotypes within each functional class (**Fig. 6**). At the group level, MMR mutants showed stronger daily synergy scores than ODR mutants yet retained higher absolute growth (**Fig. 7**), consistent with their elevated mutation supply enabling continued growth despite synergistic drug pressure. Bliss scores could not be reliably computed for late time points under meropenem-escalating regimens, as wildtype populations approached extinction at high concentrations.

**Figure 6.**
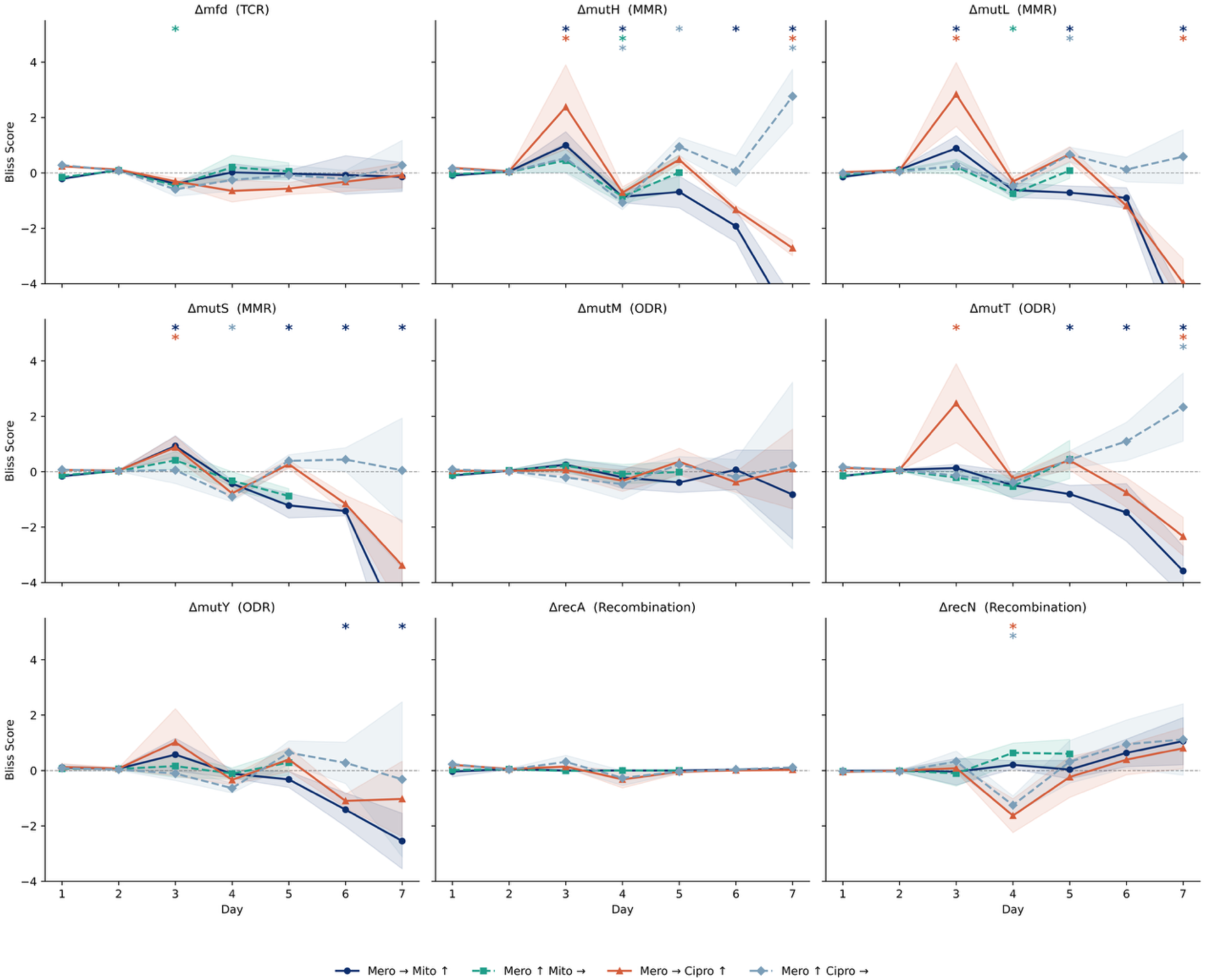
Day-resolved Bliss independence scores for individual knockout strains across four combination treatments. Each panel shows one strain; lines connect daily Bliss scores computed from 24 h OD_595 nm_ measurements, with mutant fitness defined as the ratio of mutant OD_595 nm_ to mean wildtype OD_595 nm_ under the same treatment on the same day. Shaded regions represent 95% confidence intervals. Values below zero indicate synergy. Coloured asterisks at the top of each panel mark days where the strain’s Bliss score differed significantly from the wildtype distribution (Wilcoxon rank-sum test, BH-corrected p < 0.05); colours correspond to combination treatments. Days where the mean wildtype OD_595 nm_ fell below 0.05 were excluded.

**Figure 7.**
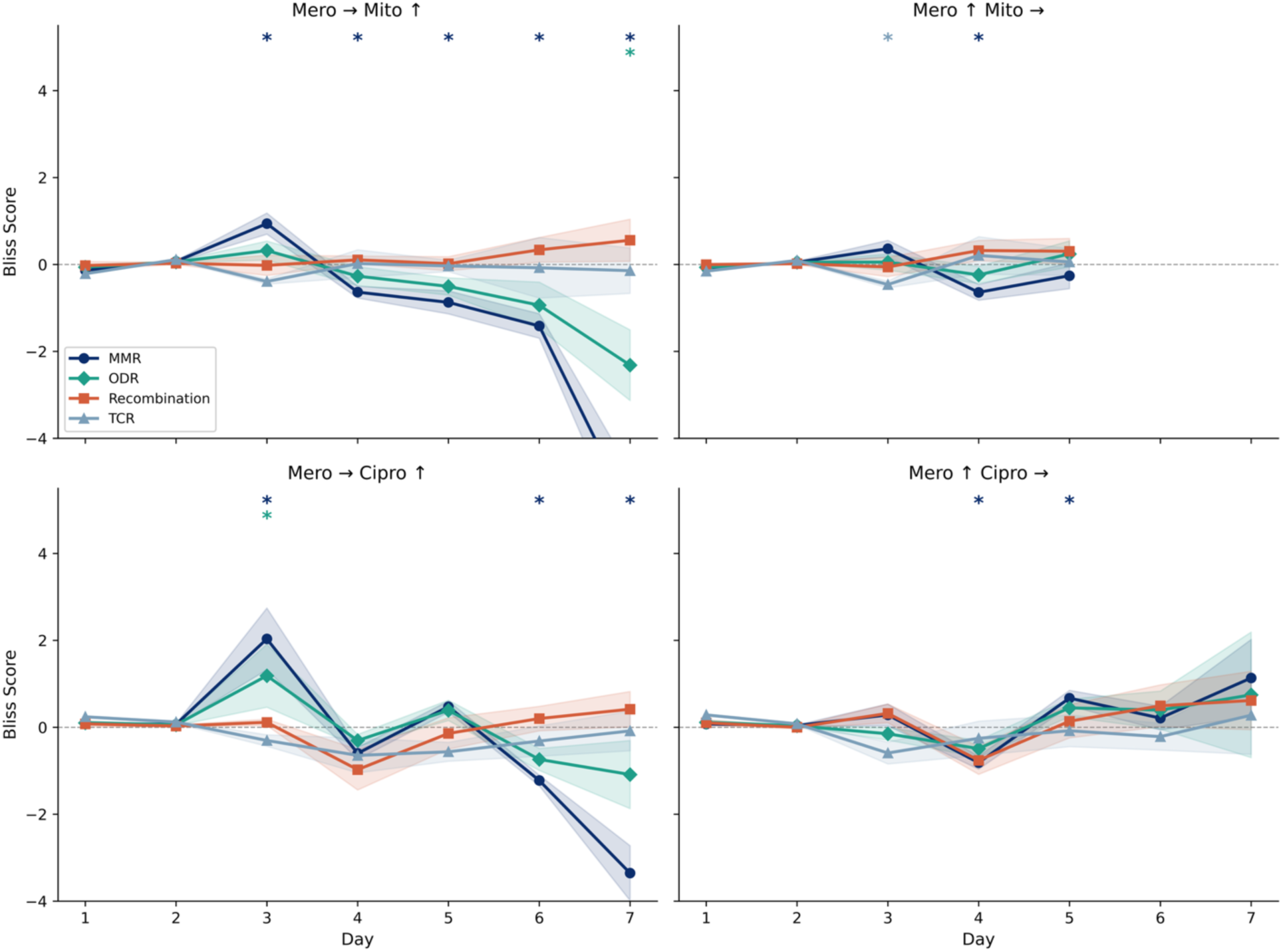
Day-resolved Bliss independence scores aggregated by functional group. MMR (Δ*mutH*, Δ*mutL*, Δ*mutS*), ODR (Δ*mutM*, Δ*mutT*, Δ*mutY*), Recombination (Δ*recA*, Δ*recN*), and TCR (Δ*mfd*) are shown for each combination treatment. Lines show the mean Bliss score per day; shaded regions represent the 95% confidence interval. Coloured asterisks at the top of each panel mark days where the group’s Bliss scores differed significantly from the wildtype distribution (Wilcoxon rank-sum test, BH-corrected p < 0.05); colours correspond to combination treatments. Days where the mean wildtype OD_595 nm_ fell below 0.05 were excluded.

## Discussion

Our results demonstrate that the principle of synthetic lethality, successfully exploited in oncology to target DNA repair-deficient tumours^9,17^, can constrain the evolvability of specific repair-deficient bacterial genotypes *in vitro*, though the degree of constraint depends on the underlying repair defect. We hypothesised that pairing antibiotic selection with DNA-damaging agents would exploit synthetic lethal interactions to selectively counteract hypermutator lineages.

The data reveal that strains deficient in double-strand break repair (Δ*recA*, Δ*recN*), transcription-coupled repair (Δ*mfd*), and oxidative damage repair (Δ*mutM*, Δ*mutT*, Δ*mutY*) were constrained under combination treatments, though the mechanistic basis for constraint likely differs across these functionally distinct groups. In contrast, classical MMR hypermutators (Δ*mutS,* Δ*mutH,* Δ*mutL*), which represent prevalent mutator genotypes in chronic clinical infections, did not show reduced evolvability under these treatments. These strains maintained greater evolvability despite the added genotoxic burden. This divergence establishes that hypermutation cannot be treated as a single phenotype; rather, the success of evolution-suppressing therapies depends on exploiting specific pathway orthogonality.

The reduced evolvability of ODR mutants supports the hypothesis that exacerbating specific genomic instabilities can constrain evolutionary trajectories. Carbapenem antibiotics, including meropenem, have been reported to induce reactive oxygen species (ROS) production as a secondary bactericidal mechanism^18–20^, although the contribution of ROS to antibiotic lethality remains debated. If operative, this mechanism would compound the vulnerability of ODR-deficient strains, which lack the enzymatic machinery to mitigate oxidative DNA damage. Bliss independence analysis confirmed that both ODR and MMR mutants experienced significant synergistic fitness costs under constant meropenem with increasing mitomycin C, and that combination treatments imposed greater fitness deficits than predicted from independent drug effects. Notably, MMR mutants showed stronger synergy than ODR mutants, yet ODR mutants exhibited substantially greater reductions in evolvability. This apparent paradox is resolved by considering the hypermutator phenotype of MMR-deficient strains (mutation rates ~100–1,000-fold higher than wildtype)^21,22^, which likely provides sufficient genetic variation for rapid adaptation despite synergistic drug pressure.

Crucially, the efficacy of this constraint was strictly modulated by the dosing regimen. For example, the regime maintaining constant antibiotic stress (via meropenem) while increasing genotoxic stress (via mitomycin C) proved more efficacious than the inverse schedule, in which antibiotic concentration was increased against a constant genotoxic background. The greater efficacy of constant meropenem with increasing mitomycin C is consistent with multiple interpretations: sustained antibiotic pressure may ensure continuous ROS-mediated sensitisation to escalating genotoxic stress^23^, or alternatively, increasing mitomycin C may potentiate meropenem killing through compounding DNA damage. Distinguishing these mechanisms will require time-resolved measurements of DNA damage and repair activity. Positive Bliss scores for Δ*recA* and Δ*recN* likely reflect a floor effect rather than true antagonism: these strains showed near-complete growth inhibition under monotherapy alone, precluding meaningful assessment of drug interactions using the Bliss framework. This is consistent with RecA’s role in both homologous recombination and the induction of the SOS response^24^.

The resilience of MMR-deficient strains highlights the limitation of using agents that induce structural DNA damage to constrain specific mutators. MMR mutants function as generalist mutators, accelerating adaptation by elevating the rate of single-nucleotide polymorphisms (SNPs) through uncorrected replication errors^22,25^. However, the MMR system is orthogonal to the repair of the structural lesions induced by the genotoxic agents. MMR proteins do not participate in resolving the inter-strand crosslinks caused by mitomycin C or the double-strand breaks induced by ciprofloxacin, which are instead processed by nucleotide excision repair and homologous recombination^13,26^. Consequently, MMR-deficient strains did not exhibit the synthetic lethality associated with these structural damages. Instead, as noted, they likely benefited from their elevated mutation supply, enabling rapid adaptation to the meropenem selection ramp. This dynamic mirrors the mutator phenotype in oncology, where MMR-deficient tumors often exhibit intrinsic tolerance to specific DNA-damaging chemotherapies (such as fluoropyrimidines) because the repair machinery required to recognise the lesions and mediate cytotoxicity is absent^27^. We speculate that constraining bacterial MMR mutators may require agents that directly penalise replication fidelity, such as nucleoside analogs or inhibitors of error-prone polymerases, rather than agents that inflict structural DNA damage; however, this hypothesis remains to be tested.

Several limitations should be considered when interpreting these results. First, we cannot distinguish whether adaptation occurred through selection on pre-existing genetic variation or *de novo* mutations arising during the experiment. MMR-deficient strains may harbour elevated standing variation due to their constitutively elevated mutation rates, potentially allowing adaptation through selection alone. Whole-genome sequencing of evolved populations would be needed to resolve this. Second, the ΔAUC metric integrates both intrinsic drug sensitivity and evolutionary adaptation over the 7-day experiment; for genotypes with strong innate sensitivity (e.g., Δ*recA*), reduced ΔAUC may largely reflect the former rather than a failure to evolve *per se*. Third, our study employed laboratory-evolved populations under defined *in vitro* conditions; clinical translation will require validating these dynamics in more complex models where metabolic heterogeneity may alter ROS production and drug uptake. Fourth, our experiments address only constitutive repair deficiencies; antibiotic exposure can also transiently elevate mutation rates through inducible stress response pathways^24,28,29^, whose contribution to evolvability under combination treatments remains unexplored. Finally, we focused on single-gene knockouts from a single genetic background. Future work must address the potential for redundancy in repair pathways, epistatic interactions between repair deficiencies, and the impact of horizontal gene transfer, which can restore repair functions in open clinical systems^30^.

## Conclusion

Collectively, these findings establish that targeting genomic instability is a viable strategy for restricting the evolvability of specific hypermutator subclasses. We demonstrate that oxidative repair-deficient bacteria show markedly reduced evolvability under combination therapies, consistent with a synergy between antibiotic-mediated stress and exogenous genotoxicity. However, the distinct resilience of MMR mutants reveals that future evolution-aware therapies must be genotype-directed. By employing diagnostic precision to match therapeutic stress to the specific molecular defects of the pathogen, we can develop rationally designed combinations that substantially constrain the evolutionary trajectories of bacterial populations.

## Supplementary Information

**Supplementary Figure 1.**
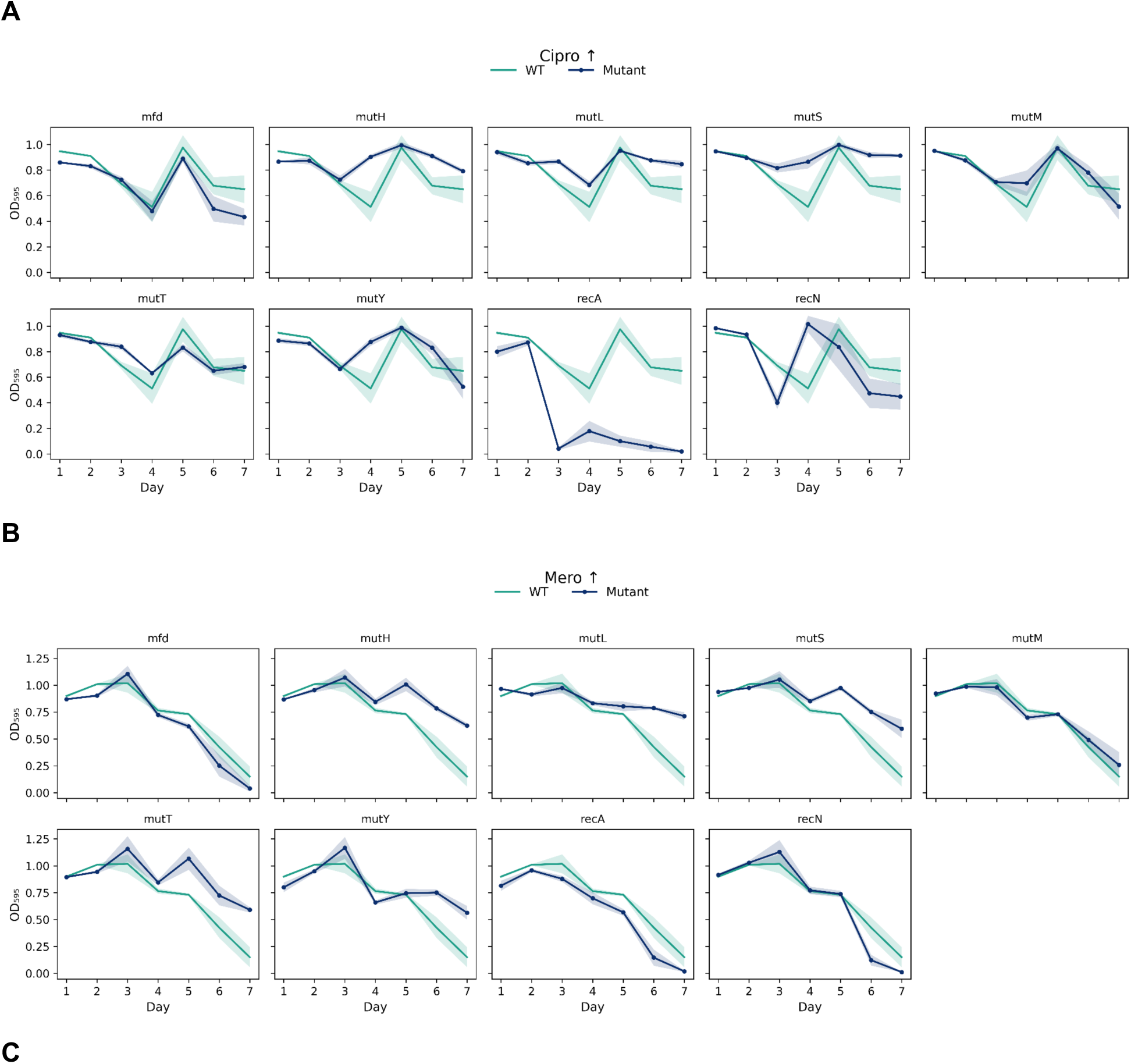
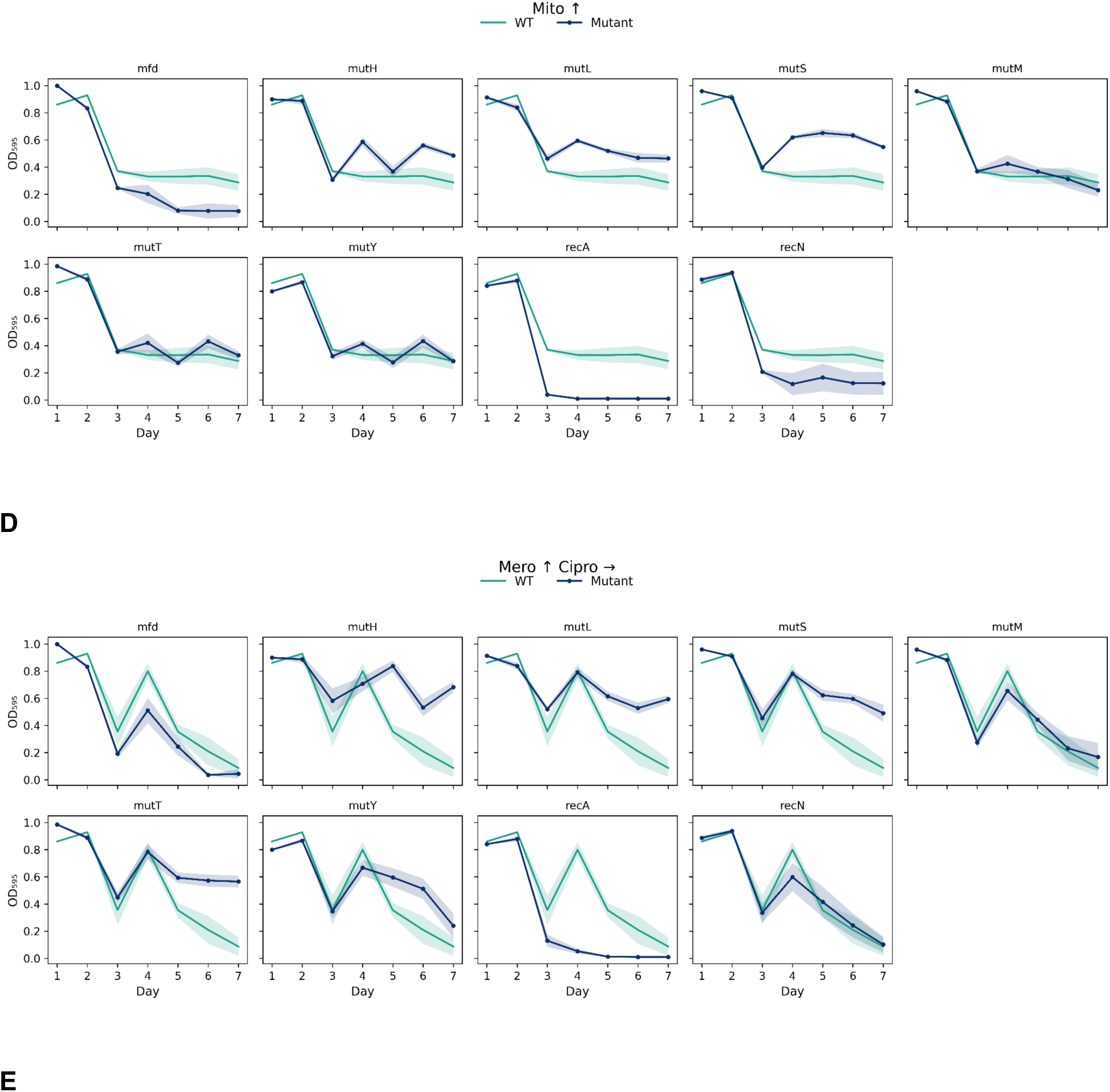
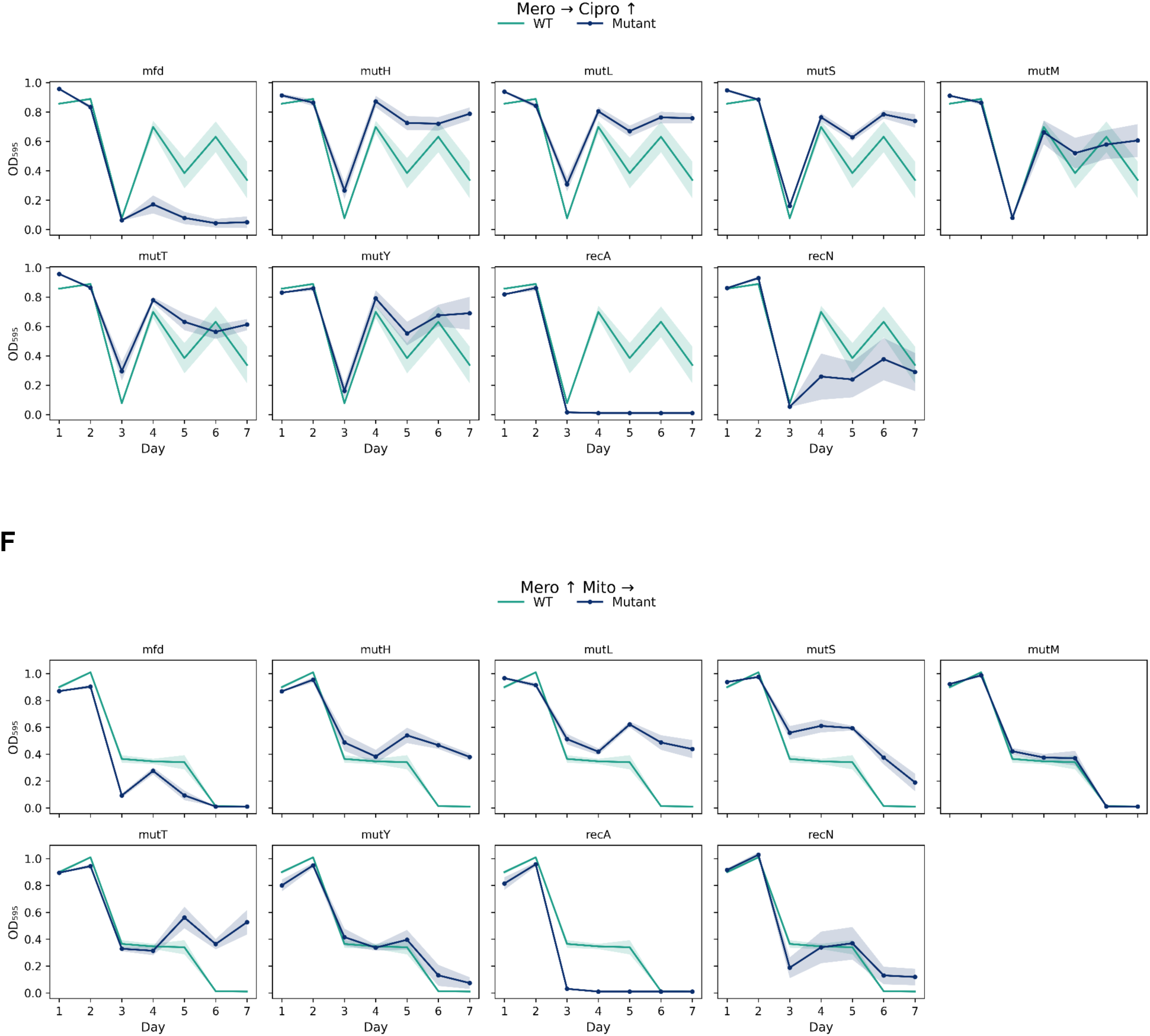
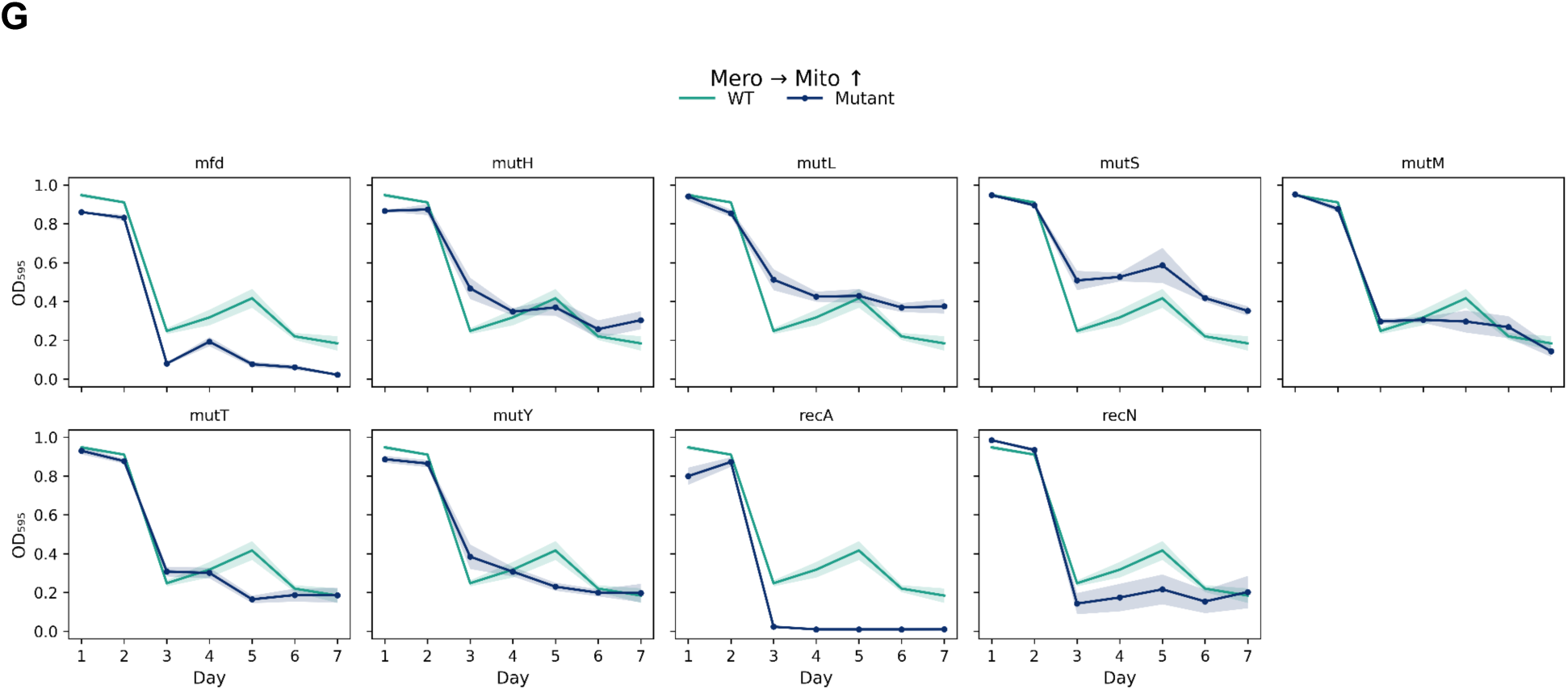
Temporal growth dynamics. Mean optical density (OD_595_) trajectories over 7 days of experimental evolution for individual single-gene knockout strains (blue lines) compared to the paired wild-type control (green lines) evolved under identical conditions. Panels are grouped by treatment regimen: (A) Ciprofloxacin escalation (Cipro ↑); (B) Meropenem escalation (Mero ↑); (C) Mitomycin C escalation (Mito ↑); (D) Meropenem escalation with constant Ciprofloxacin (Mero ↑ Cipro →); (E) Constant Meropenem with Ciprofloxacin escalation (Mero → Cipro ↑); (F) Meropenem escalation with constant Mitomycin C (Mero ↑ Mito →); (G) Constant Meropenem with Mitomycin C escalation (Mero → Mito ↑). Shaded regions represent the standard error of the mean (s.e.m.) across biological replicates.

**Supplementary Figure 2.**
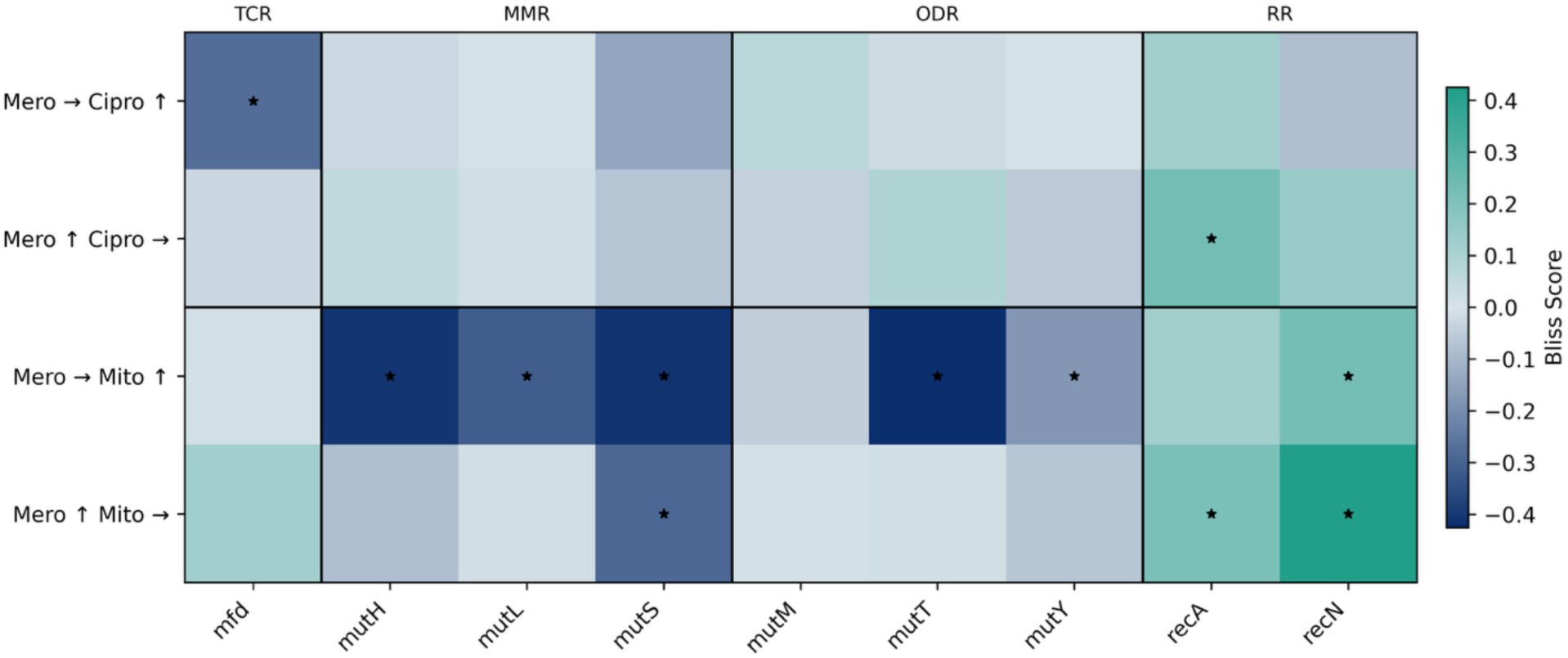
Bliss independence analysis. Heatmap showing mean Bliss scores computed from AUC values integrated across the full 7-day experiment, for each genotype–combination. Bliss score = observed combination fitness − expected fitness, where fitness is defined as mutant AUC relative to wildtype AUC. Negative values (blue) indicate synergy, where the combination imposed greater fitness costs than predicted from independent drug action; positive values (green) indicate antagonism, where the combination was less efficacious than expected. Black asterisks mark significance (Wilcoxon rank-sum test, mutant vs wildtype, Benjamini–Hochberg corrected p < 0.05). Columns are grouped by functional class. Row labels indicate combination treatment: ↑ = increasing concentration, → = constant concentration.

## Methods

### Strains and media

Single-gene knockout strains were obtained from the Escherichia coli K-12 BW25113 KEIO collection^11,12^. Strains carrying deletions in mismatch repair (Δ*mutS*, Δ*mutH*, Δ*mutL*), oxidative damage repair (Δ*mutM*, Δ*mutT*, Δ*mutY*), double-strand break repair (Δ*recA*, Δ*recN*), and transcription-coupled repair (Δ*mfd*) were selected for evolution experiments. The isogenic wildtype BW25113 served as the control. Strains were maintained on LB-agar plates containing 50 µg/mL kanamycin at 37 °C and stored at −80 °C in 20% glycerol for long-term storage. All evolution experiments were conducted in Mueller-Hinton Broth (MHB; Merck, cat. no. 70192).

### Antibiotics and DNA-damaging agents

Meropenem, ciprofloxacin, and mitomycin C were purchased from Sigma-Aldrich. Stock solutions were prepared according to the supplier’s recommendations using water as the solvent and stored at −30 °C until use. Minimum inhibitory concentrations (MICs) were determined for the wildtype strain using standard broth microdilution in MHB (meropenem = 0.125 µg/mL; ciprofloxacin = 0.016 µg/mL; mitomycin C = 1 µg/mL).

### High-throughput experimental evolution

Evolution experiments were performed in 384-deep well plates (Greiner Bio-One) using a high-throughput liquid handling platform (Biomek FXP, Beckman Coulter). Strains were streaked from glycerol stocks onto LB-agar plates containing 50 µg/mL kanamycin, grown overnight at 37 °C, and single colonies were used to inoculate 384-deep well plates containing 150 µl MHB per well (8 replicates per strain). Plates were sealed with gas-permeable membranes (AeraSeal, Sigma-Aldrich), centrifuged briefly (600 rpm, 2 min), and incubated at 37 °C with shaking at 600 rpm for 24 h before each transfer.

Populations were evolved for 7 days under a stepwise drug selection ramp. Each day, cultures were diluted 1:100 into fresh MHB containing the specified drug concentration(s). For monotherapy treatments, drug concentrations were doubled daily from 0.5× to 32× MIC. For combination treatments, one drug followed the same escalating ramp while the partner drug was maintained at a constant sub-inhibitory concentration (0.5× MIC) throughout the experiment. Both treatment orientations were tested for each drug pair (i.e., Drug A increasing while Drug B is held constant, and vice versa).

### Optical density measurements

Optical density at 595 nm (OD_595_) was measured daily prior to each transfer. Aliquots from 384-deep well plates were transferred to shallow flat-bottom 384-well microplates (Greiner Bio-One) and measured using an Infinite M1000 Pro plate reader (Tecan). Background absorbance from sterile media wells was subtracted from all measurements.

### Modelling of mutant–wildtype growth differences

Differences between mutant and wildtype growth were quantified using Bayesian modelling framework applied to AUC measurements. Mutant and wildtype observations were modelled independently, with drug-specific baselines shared across genotypes and treatment-specific shifts (ΔAUC) estimated for each mutant–drug combination. Each biological replicate (mutant or wildtype) appeared exactly once in the dataset, ensuring that the effective sample size correctly reflected the number of independent observations.

The primary quantity of interest was the posterior distribution of the mutant–treatment effect (ΔAUC), defined as the difference in AUC between mutant and wildtype under a given condition. Throughout, ΔAUC was interpreted as a proxy for realised evolvability under drug exposure, with negative values indicating stronger suppression of mutant growth relative to wildtype.

Uncertainty was quantified using posterior credible intervals, and effects were summarised using posterior means and signed tail probabilities to capture both the direction and strength of mutant responses. For functional classes of mutants, group-level effects were computed by averaging posterior draws across mutants within each class, yielding posterior distributions that correctly propagate uncertainty. Full model specification, priors, diagnostics, and posterior predictive checks are provided in the **Supplementary File 1**.

### Bliss independence analysis

To assess whether combination treatments produced non-additive fitness effects, we computed the Bliss independence score (ε) for each mutant genotype under each combination treatment. For each mutant, we calculated relative fitness as the ratio of the mutant’s AUC to wildtype mean AUC under each treatment condition. The expected fitness under combination treatment was computed as the product of relative fitnesses under the component monotherapies (Bliss independence model):

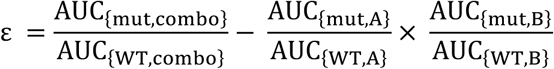

Negative Bliss scores indicate synergistic interactions in which the combination imposed greater fitness costs than predicted from the individual drug effects; positive scores indicate antagonistic interactions. To control for non-specific effects arising from differences in concentration dynamics between monotherapy and combination regimens, Bliss scores were also computed for wildtype replicates under all combination treatments. Statistical significance was assessed using two-sided Wilcoxon rank-sum tests comparing mutant Bliss scores to wildtype Bliss scores under the same combination, with Benjamini–Hochberg correction for multiple testing across all strain–combination comparisons. This approach tests whether mutant Bliss scores differ from the wildtype reference distribution, thereby controlling for any systematic deviations from additivity that arise from the experimental design rather than genotype-specific drug interactions.

To assess the temporal dynamics of drug interactions, we additionally computed day-resolved Bliss scores using daily OD measurements rather than integrated AUC values. For each day, mutant fitness was calculated as the ratio of mutant OD to the mean wildtype OD under the same treatment. Days on which mean wildtype OD fell below 0.05 under any component treatment were excluded to avoid unstable fitness ratios arising from near-zero denominators. For day-resolved analyses, significance was assessed using Wilcoxon rank-sum tests comparing daily mutant Bliss scores to wildtype Bliss scores on the same day under the same combination, with Benjamini–Hochberg correction. Significance markers in Figures 6 and 7 are restricted to Day ≥ 3 (when drug concentrations exceed the MIC) and require an absolute effect size Δbliss > 0.2, Day ≥ 3.

## Code and data availability

Raw data will be archived on Zenodo. The code will be uploaded to the GitHub repository <url> and also archived on Zenodo.

## Funding

WM was supported by the Wellcome Early-Career Award 319534/Z/24/Z. This work was supported by UKRI Frontiers Grant (EP/Y031067/1) to RCM. This work was also funded by the ERC grant uCARE (ID 819454) to A.T.. S.O.B. was supported by the Add-On Fellowship for Interdisciplinary Science from the Joachim Herz Foundation.

## Notes

### Competing Interest Statement

The authors have declared no competing interest.

## References

1. Ho, C. S. et al. Antimicrobial resistance: a concise update. Lancet Microbe 6, (2025).

2. Cox, E. C. Bacterial mutator genes and the control of spontaneous mutation. Annu. Rev. Genet. 10, 135–156 (1976).

3. Gifford, D. R. et al. Mutators can drive the evolution of multi-resistance to antibiotics. PLOS Genet. 19, e1010791 (2023).

4. Oliver, A. & Mena, A. Bacterial hypermutation in cystic fibrosis, not only for antibiotic resistance. Clin. Microbiol. Infect. 16, 798–808 (2010).

5. Pal, C., Maciá, M. D., Oliver, A., Schachar, I. & Buckling, A. Coevolution with viruses drives the evolution of bacterial mutation rates. Nature 450, 1079–1081 (2007).

6. Perron, G. G., Hall, A. R. & Buckling, A. Hypermutability and Compensatory Adaptation in Antibiotic-Resistant Bacteria. Am. Nat. 176, 303–311 (2010).

7. Campbell, B. B. et al. Comprehensive Analysis of Hypermutation in Human Cancer. Cell 171, 1042–1056.e10 (2017).

8. O’Connor, M. J. Targeting the DNA Damage Response in Cancer. Mol. Cell 60, 547–560 (2015).

9. Farmer, H. et al. Targeting the DNA repair defect in BRCA mutant cells as a therapeutic strategy. Nature 434, 917–921 (2005).

10. Holohan, C., Van Schaeybroeck, S., Longley, D. B. & Johnston, P. G. Cancer drug resistance: an evolving paradigm. Nat. Rev. Cancer 13, 714–726 (2013).

11. Baba, T. et al. Construction of Escherichia coli K-12 in-frame, single-gene knockout mutants: the Keio collection. Mol. Syst. Biol. 2, 2006.0008 (2006).

12. Yamamoto, N. et al. Update on the Keio collection of Escherichia coli single-gene deletion mutants. Mol. Syst. Biol. 5, 335 (2009).

13. Drlica, K. & Zhao, X. DNA gyrase, topoisomerase IV, and the 4-quinolones. Microbiol. Mol. Biol. Rev. MMBR 61, 377–392 (1997).

14. Iyer, V. N. & Szybalski, W. A molecular mechanism of mitomycin action: linking of complementary dna strands*†. Proc. Natl. Acad. Sci. 50, 355–362 (1963).

15. Gifford, D. R. et al. Identifying and exploiting genes that potentiate the evolution of antibiotic resistance. *Nat*. Ecol. Evol. 2, 1033–1039 (2018).

16. San Millan, A., Escudero, J. A., Gifford, D. R., Mazel, D. & MacLean, R. C. Multicopy plasmids potentiate the evolution of antibiotic resistance in bacteria. *Nat*. Ecol. Evol. 1, 1–8 (2016).

17. Lord, C. J. & Ashworth, A. PARP inhibitors: Synthetic lethality in the clinic. Science 355, 1152–1158 (2017).

18. Kohanski, M. A., Dwyer, D. J., Hayete, B., Lawrence, C. A. & Collins, J. J. A Common Mechanism of Cellular Death Induced by Bactericidal Antibiotics. Cell 130, 797–810 (2007).

19. Dwyer, D. J. et al. Antibiotics induce redox-related physiological alterations as part of their lethality. Proc. Natl. Acad. Sci. U. S. A. 111, E2100–2109 (2014).

20. Van Acker, H. & Coenye, T. The Role of Reactive Oxygen Species in Antibiotic-Mediated Killing of Bacteria. Trends Microbiol. 25, 456–466 (2017).

21. Lee, H., Popodi, E., Tang, H. & Foster, P. L. Rate and molecular spectrum of spontaneous mutations in the bacterium Escherichia coli as determined by whole-genome sequencing. Proc. Natl. Acad. Sci. 109, E2774–E2783 (2012).

22. Schofield, M. J. & Hsieh, P. DNA mismatch repair: molecular mechanisms and biological function. Annu. Rev. Microbiol. 57, 579–608 (2003).

23. Foti, J. J., Devadoss, B., Winkler, J. A., Collins, J. J. & Walker, G. C. Oxidation of the Guanine Nucleotide Pool Underlies Cell Death by Bactericidal Antibiotics. Science 336, 315–319 (2012).

24. Cirz, R. T. et al. Inhibition of Mutation and Combating the Evolution of Antibiotic Resistance. PLOS Biol. 3, e176 (2005).

25. Tenaillon, O. et al. Tempo and mode of genome evolution in a 50,000-generation experiment. Nature 536, 165–170 (2016).

26. Dronkert, M. L. G. & Kanaar, R. Repair of DNA interstrand cross-links. Mutat. Res. Repair 486, 217–247 (2001).

27. Sargent, D. J. et al. Defective Mismatch Repair As a Predictive Marker for Lack of Efficacy of Fluorouracil-Based Adjuvant Therapy in Colon Cancer. J. Clin. Oncol. 28, 3219–3226 (2010).

28. Andersson, D. I. & Hughes, D. Microbiological effects of sublethal levels of antibiotics. Nat. Rev. Microbiol. 12, 465–478 (2014).

29. Eliopoulos, G. M. & Blázquez, J. Hypermutation as a Factor Contributing to the Acquisition of Antimicrobial Resistance. Clin. Infect. Dis. 37, 1201–1209 (2003).

30. Denamur, E. et al. Evolutionary Implications of the Frequent Horizontal Transfer of Mismatch Repair Genes. Cell 103, 711–721 (2000).

